# Conditions and properties of non-Mendelian transmission of recombinants in female meiosis

**DOI:** 10.64898/2026.09.01.748656

**Authors:** Ankita Chauhan, Nathan Yeung, Helen Kim, Kevin H-C Wei

## Abstract

In female meiosis, each of the four chromatids in a tetrad has a 25% chance to be selected for transmission through the pronucleus, realizing Mendel’s second law. Here, we characterize a cheating but unselfish behaviour that has documented cases across diverse animal taxa including humans and flies, whereby chromatids with crossovers (COs) have a transmission advantage, resulting in increased production of recombinant offspring. Taking advantage of *Drosophila* ovarian physiology, we show that this form of meiotic cheating which we call recombinant drive occurs on the autosomes when females are under nutrient stress, potentially as a mechanism for recombination plasticity, while curiously, the effect is suppressed on the X by the distributive system. We explored recombinant drive using simulations and identified unique quantitative signatures that are at odds with several intrinsic properties of linkage. One violation is the production of more offspring with recombinant versus parental allele combinations, which we empirically demonstrated to be possible. Further, we show that recombinant drive can appear to modify CO patterning effectively acting as assurance and interference mechanisms, even when it has no influence on and acts downstream of CO spacing. We discuss the potential benefits and consequences of having a conserved method that can rapidly increase recombination in response to stress, and speculate on a mechanism for the preferential transmission of COs at meiosis II. Overall, our study revealed distinct properties and behaviours of a poorly understood, but potentially widespread, conserved phenomenon and offers avenues for broad detection and mechanistic dissection.

**SIGNIFICANCE:** Meiotic drivers, typically described as selfish, violate Mendel’s law of segregation to gain a transmission and evolutionary advantage. Here, we explore a form of unselfish drive, whereby chromatids with crossovers are more likely to be transmitted in female meiosis, resulting in increased production of offspring with novel allelic combinations. This is a form of Mendelian cheating called recombinant drive, and has been documented in some animals, including humans and flies. Using classical genetics and computational simulations, we uncovered unique consequences of recombinant drive, some of which even violate fundamental properties of recombination and linkage. Overall, our study begins to illuminate an otherwise poorly characterized but potentially conserved mechanism that has implications on evolutionary and population dynamics.

## INTRODUCTION

Mendel’s second law, law of segregation, manifests in equal transmission probability of the homologous, nonsister alleles. In female meiosis, which is asymmetric, this law is realized when one of the four gametic products is randomly selected for inclusion into the pronucleus to be fertilized and the remainder becomes polar bodies to be discarded. Compared to the symmetry of male meiosis which produces four viable gametes, the asymmetry of female meiosis offers a unique opportunity for violation to Mendel’s second law. In a heterozygote, if one of the two alleles can manipulate the segregation machineries causing preferential inclusion into the pronucleus, the allele, known as a meiotic driver, can outcompete its counterpart and spread in the population (1). Importantly, this does not directly impact female fitness as the reproductive output is unchanged.

Female meiotic drive has typically been discussed in the purview of selfish activity, such as selfish centromeres that gain transmission advantage irrespective of impact on host fitness with several examples in plants and animals (2–7). However, the asymmetry also opens the possibility of a manner of transmission distortion that is unselfish, with potential benefits to the host. One of the essential steps in meiosis is meiotic recombination that occurs through the physical crossovers (COs) of homologous chromosomes during prophase I (Figure 1A). Recombination allows parental alleles to shuffle, creating novel combinations that make natural selection more efficacious and in its absence, organisms are doomed to extinction due to a combination of population genetic effects including linked selection and Muller’s Ratchet (reviewed in (8–10)). Several lines of evidence have suggested that recombinant chromatids, the products of COs, can be preferentially transmitted over the non-recombinants. For example, in a E1 meiotic tetrad, i.e. tetrad with 1 CO, the two homologous nonsisters that participated in the CO have increased chance of inclusion into the pronucleus over the other two which are more likely to be relegated to the polar bodies (Figure 1A); preferential segregation then favors not any specific allele but novel, nonparental combinations of alleles. This was most directly demonstrated by Ottolini et al. (11) who examined human oocytes donated from in vitro clinics; by isolating and individually sequencing all nuclei within oocytes, they observed an excess of recombinants in the pronuclei compared to polar bodies. Although less direct, evidence of the same phenomenon has also been observed in different species of *Drosophila* (12–14). Across these cases, preferential transmission of recombinants, or recombinant drive henceforth, increases the production of recombinant offspring without the need to create more COs. This is proposed to confer an evolutionary advantage by increasing the allelic diversity of the offspring, providing a cellular mechanism for recombination rate plasticity (12, 15).

**Figure 1.**
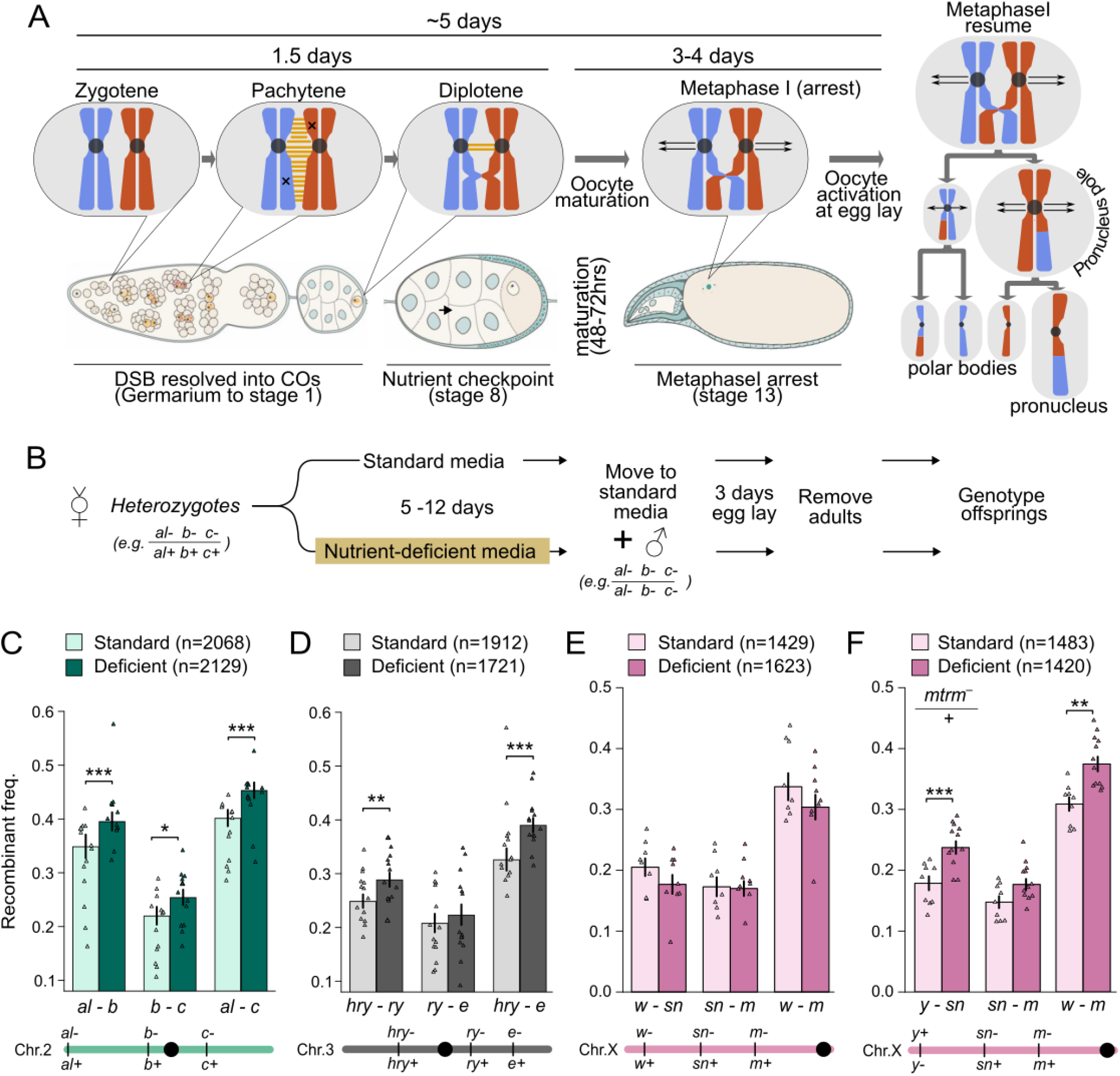
Exposing recombinant drive by timing *Drosophila* oogenesis. **A.** Meiotic progression of an E1 tetrad through in *Drosophila* oogenesis. Oocyte diagrams illustrate stages of interest and are adapted from (20) with permission. X’s indicate double-strand breaks (DSB), and yellow bars indicate the formation of the synaptonemal complex. **B.** Time course for nutrient deprivation assay to detect recombination rate changes due to recombinant drive. **C-F.** Bars indicate the recombinant frequency for intervals across chromosomes 2 (C), 3 (D), and X (E,F) for the standard, nutrient-rich (control) and nutrient-deficient conditions. Individual points represent replicates with the total number of offspring genotyped indicated by n. Error bars represent standard error. Placement of the markers along the chromosome and the genotypes of the heterozygous virgin female are represented below the barplots. For F, the female is additionally heterozygous for a mutant *mtrm* allele on 3L. *, **, and *** indicate p-values of <0.05, <0.01, and <0.005, respectively, Chi-squared test.

In addition to increasing recombinant offspring, recombinant drive has also been shown to enable the maintenance of heterozygosity in several parthenogenetic animals, including ants (16), roundworms (17), and tardigrades (18). If recombination occurs during meiosis of parthenogenetic animals, loss of heterozygosity is expected when chromatids segregate randomly. In these animals, however, the reciprocal recombinant chromatids cosegregate resulting in diploid oocytes that maintain heterozygosity despite having different haplotypes on each chromosome as compared to the parents. This then allows the species to maintain nucleotide diversity longterm, escaping inbreeding depression which is expected of asexual reproduction through automixis (19). In these cases, manipulation of segregation was not only advantageous, but likely necessary for population survival long term. Although the underlying molecular mechanism is poorly understood, the fact that distinct taxa were able to implement a similar strategy suggests that recombinant drive may be a conserved and fundamental mechanism to regulate the transmission of recombinant chromatids.

To shed light on a still poorly understood phenomenon, we took a two-pronged approach to characterize properties of recombinant drive, combining empirical investigation and *in silico* simulations. First, we devised a simple strategy to expose the effect of recombinant drive that takes advantage of features of *Drosophila* ovarian physiology and genetics (20), enabling us to demonstrate the effect of recombinant drive on the 2nd, 3rd, and X chromosomes. Second, we simulated *in silico* COs at meiotic tetrads with driving chromatids to reveal the consequences and limits on different quantitative measures of recombination and linkage. We identified unique signatures that deviate from universal features of fair meiosis, which we then empirically validated. Our results offer strategies and conditions to trigger and diagnose recombinant drive that not only enable future molecular dissection in *Drosophila* but also detection across widespread taxa.

## RESULTS

### Nutrient deprivation increases recombination via recombinant drive

Without directly visualizing preferential segregation, the challenge with detecting transmission distortion of recombinant chromatids is in differentiating it from changes in the rate of CO formation, CO rate henceforth. Previously, Singh et al. (12) took advantage of the timing of *Drosophila* oogenesis to attribute increased recombination rate after infection to recombinant drive. Because it takes ∼5 days for an oocyte undergoing COs during prophase I to reach maturity (Figure 1A) (21), stress-induced changes in CO rate can only be observed in eggs laid 5 days after infection. Then, eggs laid within 5 days must already have COs fully established before the treatment, and any recombination rate changes can be attributed to processes downstream of CO formation. Building on this clever approach, we further take advantage of another unique feature of *Drosophila* ovarian biology, where nutrient-depletion triggers a germline checkpoint that pauses the production of new oocytes and resorption of young oocytes (22). Importantly, the nutrient checkpoint occurs at mid oogenesis (stage 8) which is after diplotene when chiasmata have already been fully formed, while any oocytes that already progressed beyond can continue to mature. In virgin females, these oocytes will reach maturity and be held in the ovarian tract until mating which triggers egg lay and resumption of egg production (23).

Utilizing these features of ovarian physiology, we devised a scheme that allows us to detect the modifiers of recombination rate downstream of CO formation (Figure 1B). Specifically, we starve heterozygous virgin females carrying recessive markers on nutrient-deficient food lacking protein for multiple days which simultaneously acts as an exogenous stressor and physiological trigger for the nutrient checkpoint; this causes accumulation of mature oocytes with already resolved COs while blocking the production of new oocytes in the ovaries. They are then switched to standard nutrient-rich food with males which initiates egg laying. Embryos laid within the first 3 days are collected as they represent previously matured oocytes that had been held. Recombinant frequencies (RF) are then estimated between adult phenotypic markers and compared to those of control females kept as virgins on standard, nutrient-rich food.

We first elected to subject virgin females to a prolonged nutrient deprivation of 12 days, as longer time leads to drastic reduction in embryo hatching (23). For chromosome 2 carrying the markers *aristaless (al)*, *black (b)*, and *curved (c)*, a total of 2129 and 2068 F2 progeny were scored for the nutrient-deficient and control, nutrient-rich conditions. RFs of nutrient-deficient females were 0.395 between *al-b*, 0.25 between *b-c*, and 0.45 between *al-c*, all of which are significantly higher than the respective RFs in the control condition (Figure 1C; p < 0.05, chi-squared test), with an average increase of 13%. On chromosome 3 carrying the markers *hairy (hry)*, *rosy (ry)*, and *ebony (e)*, a total of 1721 and 1912 progeny were scored in the two conditions. RFs in the nutrient-deficient females were 0.29 between *hry-ry*, 0.22 between *ry-e*, and 0.39 between *hry-e*, which are all increased compared to the control condition although only *hry-ry* and *hry-e* are significant (Figure 1D; p < 0.01, chi-squared test); the intervals increased on average by 12%, closely matching the magnitude on Chr. 2. However, the X chromosome which carries the markers *white* (*w*), *singed* (*sn*), and *miniature* (*m*) showed no significant increase in RF across any interval (Figure 1E; p > 0.05, chi-squared test).

Because 12-day nutrient deprivation is associated with some degree of embryonic lethality due to declining oocyte quality (23), we considered the possibility that increased recombination rate resulted from preferential lethality of nonrecombinants. To test this possibility, we evaluated the impact our nutrient deprivation scheme has on embryo hatch rate with recombining and achiasmate chromosomes using 2nd and 3rd chromosome balancers. For both chromosomes, progeny hatch rates from balancer-carrying mothers were less impacted than those from recombining (non-balancer carrying) mothers (Figure S1). This indicates that nonrecombinant progeny are not disproportionately dying due to reduced hatching – it may be the opposite – and thus, differential hatching cannot account for the observed increases in RF. Further, we tested for changes in RF with only 5 days of nutrient deprivation, which causes no appreciable embryonic lethality (23). RF, similarly, increases across all intervals on the autosomes, though not all are significant (Figure S2), which rules out disproportionate death of nonrecombinants as the cause of increased RF.

Another source of increased RF could be mitotic recombination that repairs stress-induced double-strand breaks in the germline (22, 24), especially in held oocytes that crossed the checkpoint. Because mitotic recombination does not exhibit CO interference (25), increasing RF should be accompanied with decreasing interference, which we did not observe (see below). Thus, mitotic recombination is unlikely to be involved. To directly test this possibility, we examined Chr. 4, which is achiasmate in *Drosophila* but can produce rare recombinants through mitotic rather than meiotic recombination. We subjected heterozygous virgin females with the Chr. 4 markers *cubitus interruptus* (*ci*), *shaven* (*sv*), and *eyeless* (*ey*) to 12-day nutrient deprivation, and found no recombinant offspring in the standard and nutrient deprivation conditions (Table S1). Separately, we used a Chr. 4 marked with only *ci* and *sv* and found very low rates of recombinants in both conditions (0.002 standard vs. 0.004 nutrient-deficient; Tables S1), with no significant difference (p = 0.83, chi-squared test). Since the rare recombinants were all *ci^+^ sv^−^*individuals with no reciprocal class (*ci^−^ sv^+^*), these are likely nonrecombinant *ci^−^* individuals with incomplete penetrance of the mutant phenotype (26). The observed increases in RF, therefore, cannot be attributed to stress-induced mitotic recombination.

Having ruled out several other potential sources of increased recombination rate due to mechanisms downstream of CO formation, we conclude that the increased RF is likely the result of recombinant drive. We further tested whether this effect is consistent across genetic backgrounds by repeating the experiments using F1 females derived from a different wildtype strain (DGRP-360) and assessed RFs on Chr. 2 after 5 and 12 days of nutrient deprivation. Significant increases were observed across some but not all intervals (Figure S3), suggesting that there is strain-to-strain variation in response to stress.

### Matrimony prevents recombinant drive on the X

Because the X chromosome is freely recombining much like the autosomes, we were puzzled by the lack of recombinant drive. For reasons still not fully understood, the *Drosophila* X shows many meiotic peculiarities, including elevated rates of non-disjunction and achiasmate segregation (27). This is thought to be related to the distributive system which ensures the fidelity of achiasmate disjunction such as segregation of Chr. 4 or chromosomes with inversions (28), but the recombining X, curiously, also engages this system. Recently, the distributive system has been proposed to suppress meiotic drive, as mutation in the gene *matrimony (mtrm)*, a polo kinase inhibitor necessary for sister cohesion during achiasmate disjunction (29), causes accumulation of supernumerary B chromosomes through biased segregation (30).To test the possibility that the distributive system may also act to protect other forms of non-Mendelian segregation on the X, we introduced a dominant *mtrm* mutant allele (29) into females heterozygous for X chromosome markers. We found that RFs between *y*-*sn* and *y*-*m* were significantly higher in nutrient-deficient females (0.24 and 0.37, respectively) than in nutrient-rich females (Figure 1F; p = 0.002 and 0.0097, chi-squared test). RF between *sn*-*m* was also higher under nutrient deficiency (0.18 vs. 0.15), though borderlining significance (p = 0.08, chi-squared test). These results suggest that *mtrm* and the distributive system are involved in protecting the X chromosome from recombinant drive.

### Properties of recombinant drive without CO interference

To evaluate the magnitude to which recombinant drive can modify measures of recombination, we constructed a simulator to model COs in meiotic tetrads and implemented selection of chromatids (see Materials and Methods). Briefly, a chromosome is represented by small bins with adjustable CO rates and bins are randomized for Bernoulli draws without replacement. Successful CO at a bin is then randomly assigned to one of the four chromatids for reciprocal exchange with one of the two non-sisters. One out of the four chromatids is then selected for transmission emulating female meiosis. We devised two selection regimes to model recombinant drive representing possible extremes of how COs could be preferentially transmitted: the first is for the 0 CO chromatids to be disfavored with the rest having equal chances of transmission and the second is for the chromatid with the most COs to be preferred. We refer to the two regimes as “loser-fall” and “winner-take-all”, respectively, and evaluated how they influence CO distribution, genetic distance (the average number of COs between two loci measured in Morgan (M)), and recombination frequency (the probability of observable recombinants of two loci) behaves in response.

We first simulated a chromosome 1 Morgan (M) in length without interference which entailed a chromosome comprised of 1000 bins with uniform probability of CO at a rate of 0.002 COs per bin; CO rate will then be two COs per tetrad on average and with fair meiosis, i.e. random selection, transmitted chromatids will have one CO on average, half the amount of COs at the tetrad (31). Simulating 100,000 tetrads with fair meiosis, the distribution of COs then follows a Poisson distribution as expected (32), where 36.89% and 36.71% of the chromatids have zero and one CO, respectively (Figure 2A). In both drive regimes, the distribution shifts right with the 0CO class reduced to 13.35% reflecting the amount of E0 tetrads (i.e. tetrads with zero COs) where no chromatids can have an advantage. 1CO becomes the most frequent class as E1 tetrads will always transmit 1CO chromatids. Winner-take-all has a longer tail towards higher COs as expected. Overall, the genetic distance is increased from 1M to 1.32M and 1.68M in the two respective regimes, despite identical CO rate (Figure 2A and B).

**Figure 2.**
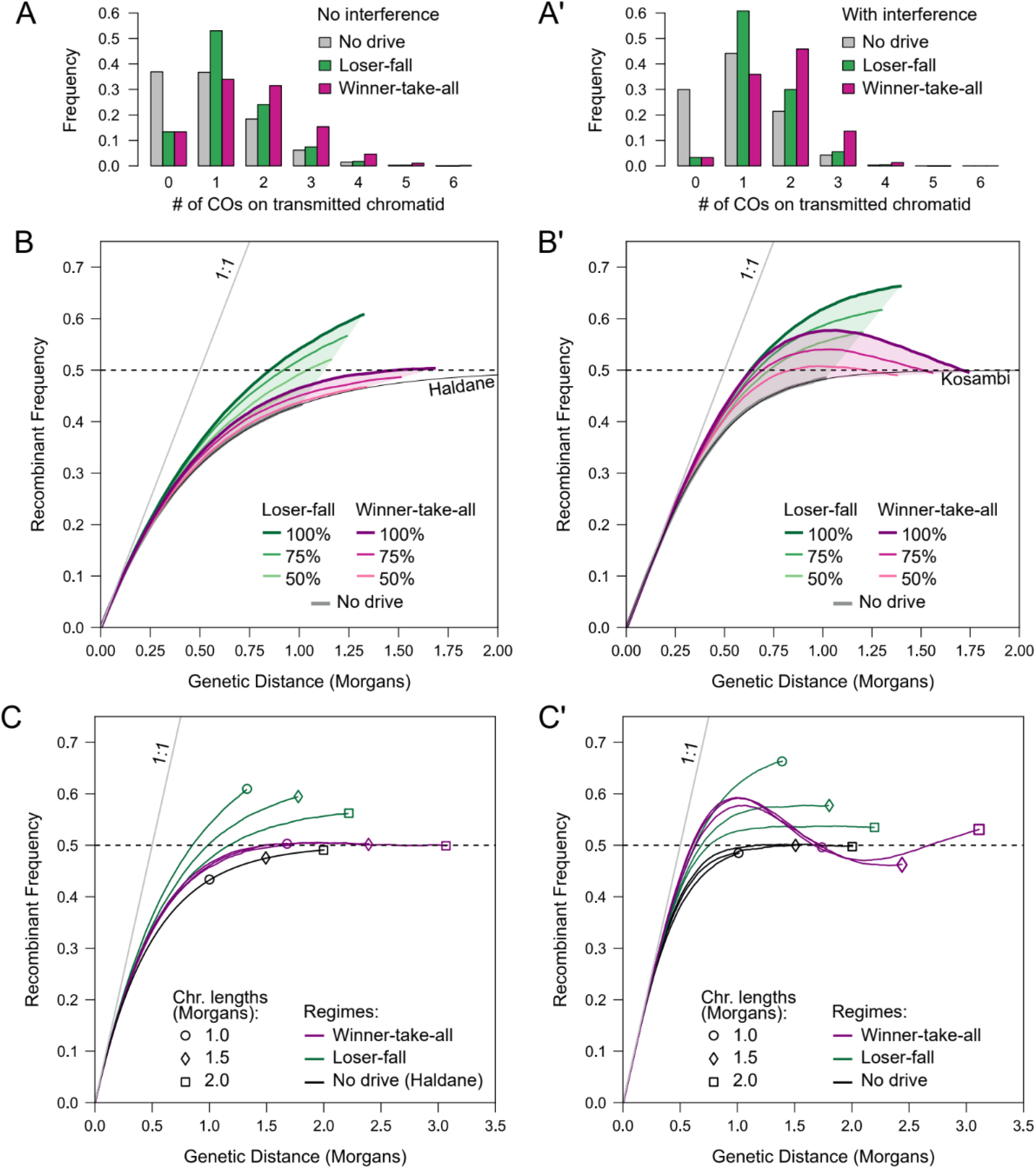
*In silico* simulations of meiotic tetrads and recombinant drive. **A.** Distribution of COs on 100,000 transmitted chromatids from 100,000 tetrads for a chromosome 1M in length. Chromatid selection from tetrads is implemented as fair (“no drive”), 0CO chromatid eliminated (“loser-fall”), or highest CO transmitted (“winner-take-all”). **B**. The relationship between genetic distance and recombinant frequency of the two drive regimes at different strengths for a 1M chromosome. Drive strength in percentage represents the probability for a tetrad to drive. Shaded areas cover the space in which genetic distance and RF can fall in the two regimes with drive strength between 0 and 100%. The Haldan map function and identity lines are plotted for reference. **C.** As with B, but with chromosomes of different lengths labeled at the termini with different symbols all driving at 100%. **A’-C’.** Same as A-C, but with interference modeled after Kosambi’s map function.

When meiosis is fair, RF asymptotes toward 50% as genetic distance increases, and when interference is absent, the relationship follows the Haldane map function (32). This is the basis of the adage that recombinants never exceed non-recombinants or parentals, although sometimes incorrectly construed as genetic distance having a maximum of 0.5M. This asymptotic relationship is partly due to the obscuring effects of double, or even number, COs which appear as non-recombinants when only two loci are considered. We examined how the increased genetic distance translates to RF with recombinant drive, and found that RF increases at a rate faster than Haldane, consistent with the distributions no longer being Poisson (Figure 2B). For winner-take-all, RF remains below, or near, 50%, maintaining the semblance of an asymptotic relationship. Surprisingly, in the loser-fall regime, despite smaller increase in genetic distance, RF exceeds 50%, reaching a terminal RF (i.e. RF between the two ends) of 60.8%. Further varying the strength of drive by adjusting the proportion of tetrads affected, we found RF capable of crossing the 50% threshold across a wide-range of conditions. This is due to an excess of 1COs chromatids (Figure 2A) which increases RF linearly with genetic distance (Figure S4A). Importantly, this is a unique signature of recombinant drive. In fair meiosis, no amount of increase in CO rate could produce RF that exceeds 0.5 (33, 34).

In addition to 1M chromosomes, we evaluated others with lengths of 1.5M and 2M (Figure 2C). Without drive, the Haldane map function is strictly followed regardless of chromosome length. In the winner-take-all regime, genetic distances of 1.5M and 2M chromosomes increased to 2.39M and 3.07M, respectively, while the terminal RF plateaus near 50%, consistent with nearly equal numbers of even and odd CO chromatids. With loser-fall, shorter chromosomes show not only steeper increases in RF, but also reach higher terminal RF. For example, the genetic distance of the 1M chromosome increased by 1.32-fold, and the terminal RF increased from 43.5% to 60.8%, a 1.40-fold increase; in contrast, the genetic distance of the 2M chromosome increased by only 1.11-fold and the terminal RF increased from 49.1% to 56.2%, a moderate 1.14-fold increase. This is because shorter chromosomes are more likely to have E1 tetrads as the most abundant recombinant configuration and they will always produce 1CO chromatids that increase RF, whereas longer chromosomes will have tetrads with more COs (e.g. E2, E3, etc), each of which can lead to transmission of both even and odd chromatids, thus buffering the increase in RF.

### Properties of recombinant drive with interference

We further implemented CO interference by depressing the CO rate in bins nearby one with a successful draw. We specifically modeled interference based on Kosambi’s formulation (35), where interference decreases proportionate to genetic distance (see Materials and Methods). For a chromosome of 1M with fair meiosis, i.e. transmitted chromatids have one CO on average, our implementation of interference increases the proportion of 1CO chromatids, while reducing those carrying larger numbers of COs compared to tetrads with no interference (Figure 2A’). Upon drive, the distribution shifts rightward to more COs, increasing the genetic distance to 1.39M, and 1.74M for the loser-fall and winner-take-all regimes, respectively. Compared to no interference, the greater impact to genetic distance here is due to fewer E0 tetrads which are insensitive to the effect of recombinant drive. Therefore, in the presence of interference, genetic distance can be further stretched by recombinant drive.

As 1CO chromatids become more numerous with interference, RF of the loser-fall regime is further exaggerated with a steeper rise and reaches a peak, terminal value of 66.2%. Strikingly, the winner-take-all regime produces a pattern with RF peaking at 57.8% when genetic distance reaches 1.06M and, curiously, decreasing after. This violates the expectation that RF increases monotonically with genetic distance in fair meiosis (32–34). The reason is that interference spaces COs further apart, causing placement of COs in 2CO chromatids, the predominant class, to be concentrated near chromosome ends. For these 2CO chromatids, the RF starting from one end of the chromosome will then increase at the beginning due to the first of the two COs and then decrease when the second of the two COs becomes abundant near the distal end, creating a negative parabolic shape (Figure S4B).

Further considering chromosomes of different lengths, we find that the loser-fall regime shows a pattern similar to that observed with no interference where the shorter chromosomes have more drastic changes. The winner-take-all regime, on the other hand, extends the streak of striking patterns where the RF not only can decrease, but, on longer chromosomes, fluctuates between decreasing and increasing which is due to increasing number of evenly spaced 3CO chromatids creating a cubic function (Figure S4B). The periodicity is expected to further continue with even longer chromosomes and can change depending on the strength and spread of interference.

### Recombination frequency exceeds 50% upon drive

Our simulations revealed that RF can exceed 50%, causing more recombinants than parentals provided that genetic distance is sufficiently long. This is a clear violation to the fundamental expectation that recombinants do not exceed parentals. We sought to test this prediction by taking advantage of the second chromosome which is ∼1.1M and has visible markers near the ends that are over 1M apart, a distance sufficient for RF to exceed the 50% threshold across most drive conditions (Figure 3A). The RFs between the markers *al* and *bw* (∼1.02M) and *al* and *speck* (∼1.06M) are 0.461 and 0.457, respectively, in the control, nutrient-rich condition. However, upon nutrient deprivation, they increased to 0.534 and 0.548, which are both subtle but significant departures from 0.5 (p = 0.03 and 0.004, Binomial exact test). Similar significant deviations were also observed in the DGRP-360 background instead of Oregon-R, where the RF increased to 0.526 and 0.531, respectively (p = 0.01 and 0.003, Binomial exact test).

**Figure 3.**
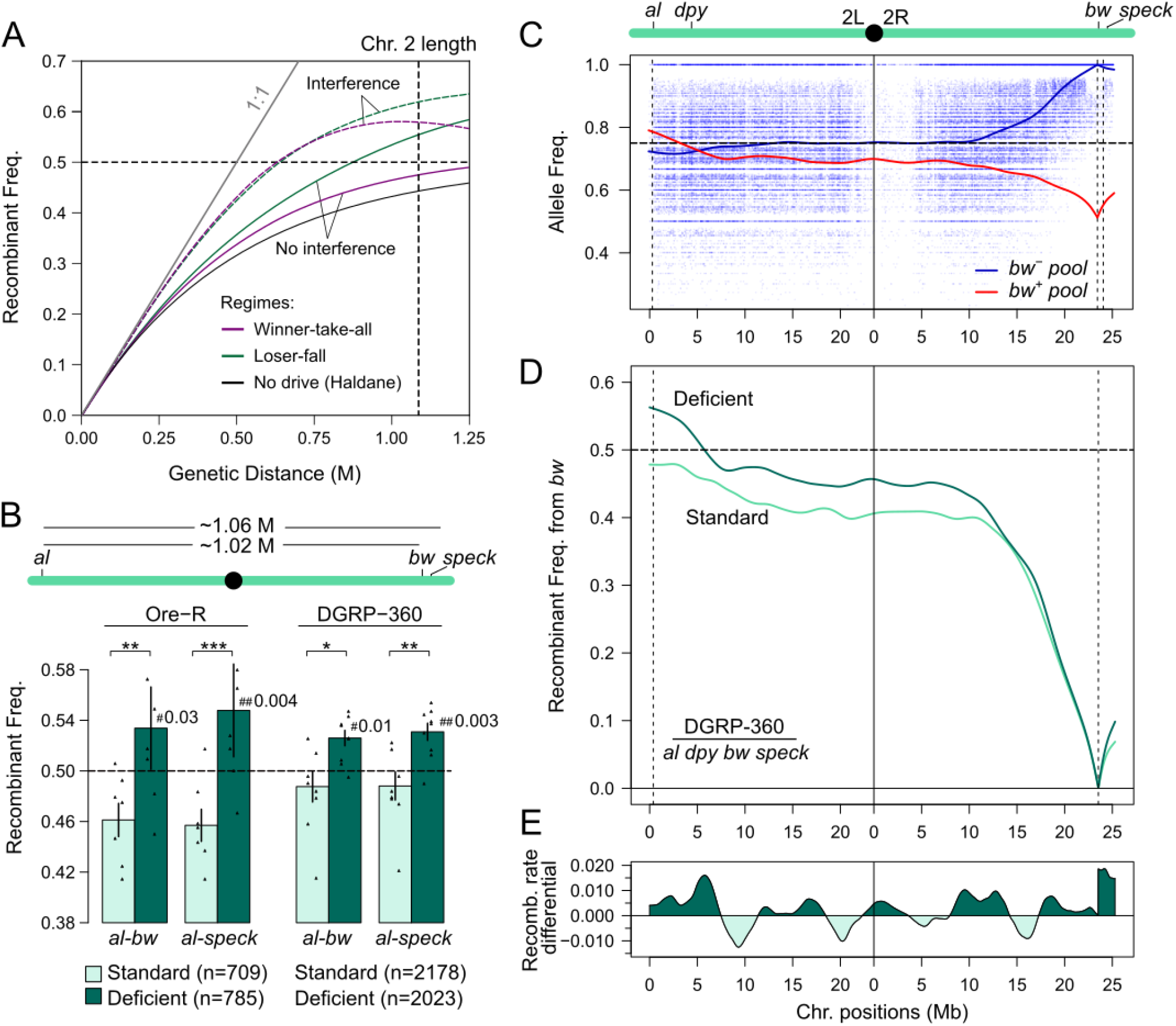
Validation of recombinant frequency exceeding 0.5 with drive. **A.** Simulation of the effect of drive on a chromosome the same length as Chr. 2 in Drosophila (1.1M). **B.** RF between intervals of the furthest markers on chromosome 2 as shown in schematic at the top. *, **, and *** indicate p-values of <0.05, <0.01, and <0.005, respectively, when comparing the RF between the conditions with the chi-squared test. # indicates the p-values from testing whether RF significantly deviates from 0.5 with the binomial exact test. **C.** Chromosome-wide allele frequency of the two conditions is estimated using marker-selected pools. For each condition, *bw^−^* and *bw^+^* pools are collected and sequenced. For illustrative purposes, allele frequency estimates of the two marker-selected pools (blue for the *bw^−^* and red for the *bw^+^* pool, respectively) are shown for the nutrient-deficient condition. Blue points represent allele frequency of the *bw^−^* pool at individual diagnostic SNP positions. The dotted vertical lines indicate scored markers and the dotted horizontal line indicates Mendelian ratio of 0.75. **D.** RF estimates based on allele frequency of the pools shown in C. **E.** The slopes of the RF curves in D were subtracted between the conditions to determine the differential. Dark and light areas indicate regions with higher and lower recombination rate due to drive in the nutrient-deficient condition, respectively.

For a chromosome-wide view of the effect of recombinant drive, we applied an approach that measures chromosome-wide RF called marker-selected pools (36). In short, when individuals from a recombinant testcross are pooled based on a selected marker, the allele frequency surrounding the marker will attenuate towards Mendelian ratio at a rate proportional to RF which can be estimated when the pool is whole-genome-sequenced. This method has the added benefit of eliminating the possibility of genotyping error for markers with more subtle phenotypes or incomplete penetrance as could be the case for *speck* and *al*. We collected and sequenced *bw^+^* and *bw^−^* pools from the two conditions and inferred the chromosome-wide allele frequency at diagnostic SNP sites (Figure 3C) which is then converted to RF (Figure 3D). Consistent with the scoring results, RF increased with nutrient deficiency and is elevated across the entire chromosome, reaching a maximum of 0.56 at the telomeric end of 2L.

Notably, the two ends of the chromosome show the largest departure in RF, suggesting that COs near the ends may be better drivers. Consistently, the differential in slope of RF, which approximates recombination rate, between the two conditions fluctuates across the chromosome, but is greatest near the ends.

### Recombinant drive depresses coefficient of coincidence

CO interference manifests as the deficiency of chromosomes with more than one CO compared to the expected probability and is captured by the coefficient of coincidence (CoC) calculated as the ratio of observed over expected 2COs. Although the underlying mechanism is still debated (37), repulsion or redistribution of COs causes them to be further apart and less likely to co-occur on a chromosome than would be expected if COs are independent. Since recombinant drive acts downstream of CO formation, interference should be unaffected. However, we noticed that while no single comparison showed significant change, CoC tends to be lower in the nutrient-deficient compared to control conditions (Table S2) and across all comparisons there is a moderate but significant trend for the depression of CoC upon recombinant drive (Figure 4A; p = 0.017, paired Wilcoxon signed-rank test).

**Figure 4.**
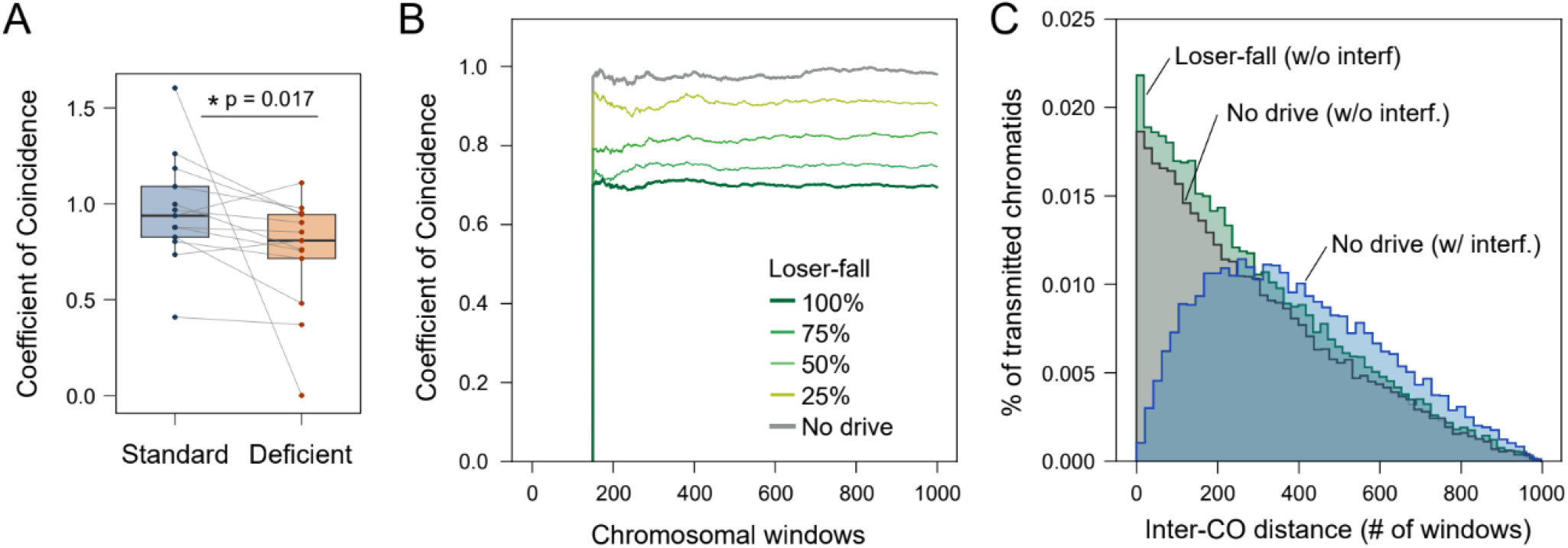
Behaviour of coefficient of coincidence with drive. **A.** Distribution of CoC estimates from all driving crosses shown in the study. Each pair of connected points represents one of the 13 experiments reported where RF was measured and compared between standard and nutrient-deficient conditions (Table S3). P-value was determined using the paired Wilcoxon signed-rank test. **B.** Based on the simulations with different strengths of drive with no interference, the CoC was calculated between pairs of chromosome windows consisting of 75 bins. The first window containing bins 1-75 was arbitrarily chosen as the reference. RF of the reference and an expanding window across the rest of the chromosome were determined and their pairwise product is then the expected rate of 2COs while the observed number is tabulated. **C.** Distribution of inter-CO distance in driving (loser-fall regime) and non-driving conditions with and without interference.

We examined this counterintuitive observation by evaluating CoC in simulations with no interference, where COs are randomly distributed in tetrads in accordance to a Poisson process. CoC was then calculated as the observed frequency of double COs between pairs of chromosome windows compared to the expected frequency calculated by the product of RFs (36, 38). The same reduction in CoC was recapitulated in the simulations (Figure 4B), affirming our empirical observation. However, there is no shift in the distribution of inter-CO distance consistent with the expectation of no change in CO repulsion (Figure 4C). The decrease in CoC, thus, does not reflect increased interference of CO formation. Rather, it is because while 1CO and 2CO chromatids both increase upon drive (Figure 2A), the increase in the former outpaces the increase in the latter, which then disproportionately increases the denominator in the CoC calculation. Therefore recombinant drive causes CoC to decrease indicating fewer chromatids with multiple COs compared to the random expectation, thus mimicking increased interference.

## yDISCUSSION

### Recombinant drive diversifies offspring as a response to stress

Here we showed that nutrient deprivation increases recombination rate through recombinant drive. This is similar to the effect of infection from parasitoid wasps and bacteria found by Singh et al (12), suggesting that recombinant drive is likely a common response to stress and may apply to many conditions that modulate recombination including temperature (39) and age (40). Towards the goal of creating more allelic combinations in the offspring, recombinant drive has several advantages over increasing CO rate. For a meaningful response to environmental stress, recombinant drive offers an immediacy that cannot be matched. As previously mentioned, the amount of time for an oocyte with new COs to become ready for fertilization is ∼5 days, which is a significant lag from the point of stress, one that may offer little benefit for short-lived organisms like flies. Changing CO rate is even less effective in many eutherian mammals including humans as COs are resolved in utero followed by prolonged late prophase I arrest (41). Then, environmental cues experienced by a female individual never impact the CO rate in her germline. Any environment-dependent mechanism to modulate offspring’ recombinant frequencies, including recombinant drive or oocyte sabotaging (42), must thus act downstream of CO formation. Moreover, even if the temporal limitation is eliminated for increasing CO rate, recombinant drive still has the advantage of higher genome-wide genetic shuffling (43) both due to the elimination of 0 CO chromatids from E1+ tetrads, as well the possibility of producing more recombinants than parentals. Lastly, recombinant drive circumvents a potential deleterious effect of increasing CO rate which also risks elevating the probability of ectopic exchange at repetitive sequences (44).

More fundamentally, the question of whether recombination rate plasticity can be under selection has been a source of much debate (45, 46). While the idea of increasing recombinant offspring upon environmental change being adaptive is intuitive (15), empirical support is limited beyond correlations. Some have attributed changes and plasticity in recombination rate as pleiotropic effects of adaptation on other features of meiosis (47). Theoretical approaches have also been mixed in their conclusions with some reporting restrictive conditions for recombination rate plasticity to be beneficial in diploid systems (48) and others showing the likely fixation of a plastic modifier locus due to associations with more favourable allele combinations (49). The presence of a stress-induced recombinant drive mechanism, which may be conserved across animals, lends support to the adaptive potential of recombination plasticity. However, this mechanism could also have emerged as a means to facilitate or stabilize chromosome disjunction and the increased recombination rate is the pleiotropic side effect. With the recombination machinery frequently under positive selection in flies (50, 51) often due to adaptation to local environments (52), how and whether recombinant drive contributes to meiotic innovations and recombination rate divergence will be of future interest.

### Mechanism of biased segregation of recombinants

Given that recombinant drive has been observed in humans and flies, it is reasonable to suppose a conserved mechanism with a common origin, which we will speculate on below. This mechanism would also provide the necessary machinery to be co-opted for the independent emergence of cosegregation of recombinant chromatids in parthenogenetic animals. For recombinants to drive, chromatid selection must occur during sister disjunction at MII since the reciprocity of COs necessitates the non-sister homologs to have equal number of COs at MI. This is consistent with the involvement of *mtrm* which ensures sister cohesion, the reduction of which may weaken the tension needed for equitable disjunction. MII drive differs from most examples of meiotic drive, particularly those relating to centromere strength (6, 42, 53–55), which occur at MI resulting in the biased transmission of the preferred allele. However, the selfish knob locus in maize drives in MII as its distance from the centromere entails high likelihood of an intervening CO that introduces the driver to the homologous chromosome creating a heterozygous sister that then drives in MII (2). The knob creates a neocentromere that has faster poleward movement (56) but does not recruit the canonical centromere architecture (57). Inspired by the knob, we envision that upon environmental stress COs can somehow form neocentromeres which then confer a stronger pull towards the pronucleus pole at MII. A scar, or “memory” (16), of COs that formed in MI must be recognizable in MII. This could occur through remnants of the CO resolution or DSB repair machinery. Since the centromeric histone CENP-A is recruited to DSB and mediates DNA repair, although in somatic human cell lines (58), the simplest scenario could be for centromeric proteins to be directly localized to COs. Alternatively, meiosis can progress in an inverted fashion with sister disjunction first followed by homolog disjunction (11, 17, 59). With inverted meiosis, resolution of COs can act to drive immediately, although the prevalence of this is currently unknown in *Drosophila*, and unlikely to be common. Resolution of the cellular mechanisms will require detailed cytological dissection of chromosomal behaviour leading to the completion of meiosis in *Drosophila.* However, some technical challenges must be overcome first as meiosis is arrested at metaphase I until egg activation and completes within a brief 5-10 minute window after egg lay (60, 61).

### Unique signatures of recombinant drive

Our simulations of recombinant drive revealed several unique signatures. The first is that RF can exceed 50% causing more recombinants than parentals provided that genetic distance is sufficiently long. The second is that with interference, RF can fluctuate between increasing and decreasing, when it is otherwise expected to increase monotonically with genetic distance. Neither of these observations, the first of which we empirically confirmed, are expected to occur with fair meiosis, thus offering diagnostic patterns to distinguish the effect of drive from CO rate increases. This can be especially useful when there is no feasible way of exploiting ovarian physiology to differentiate the sources of increasing recombination rate as was possible here with *Drosophila*.

Another interesting facet of recombinant drive revealed by our simulations is that it can, prima facie, mediate CO patterning despite acting downstream of known mechanisms (37, 62, 63). The preference for chromatids with COs in the meiotic tetrads, especially E1s, effectively acts as a mechanism for CO assurance by decreasing the number of 0CO chromatids. More puzzling, we showed the counterintuitive results that CoC decreases with drive indicating increase in interference, even when there’s no repulsion between COs at formation. The ratio of observed over expected 2COs as measured by CoC, therefore, captures mechanisms of interference beyond repulsion or spacing models (37) that are typically envisioned.

Recombinant drive, thus, has a plethora of effects on how recombination manifests, some expected, some curious, and others rule-breaking. Through this study, we have begun to reveal some of these quirks and features that we hope to be further extrapolated, dissected, and implemented in future studies.

## MATERIALS AND METHODS

### Fly handling and crosses

All *Drosophila melanogaster* strains used in this study are listed in Table S3. Flies were maintained on standard Bloomington food at 25°C with a 12:12 hour light:dark cycle. The nutrient-deficient diet is made of 44g agar, 180mL molasses, 37mL 5% Tegosept, mixed with 1112mL water followed by heating and sterilizing in the autoclave.

### Recombinant Frequency Assay

To generate heterozygous virgins, wild-type females (OreR, DGRP-360) were mated to males carrying recessive phenotypic markers on chromosomes X, 2, and 3, at 25°C on standard Bloomington food. 3-5 days old virgin F1 females were collected and transferred to vials with either the standard, nutrient-rich food or nutrient-deficient food, starting the nutrient deprivation time courses. Each replicate vial contains 10-20 females that are transferred to fresh media every 3-5 days. After the prescribed deprivation times (5-12 days, depending on experiment), the females were transferred to vials with standard Bloomington food and 10 males that have the appropriate recessive mutant genotypes; this triggers egg lay and re-initiates oogenesis. Flies were transferred to fresh vials daily to new vials for 3 consecutive days to allow for egg laying. Progeny from each daily collection were phenotypically scored for visible markers to determine recombination frequencies between marker pairs on each chromosome. Recombinant frequency was calculated as: Recombinant frequency = Number of recombinants between marker pair/ Total offspring. To introduce *mtrm*, we crossed BL147 females (*w sn m*) with BL607084 males (*yw; mtrm[exc13]/TM3-sb*). F1 heterozygous virgins were selected for the absence of TM3-sb and then placed on the different food. Wildtype males were added upon transfer to standard food, and RF were measured from the resulting euploid sons only.

### Hatchability Rate Assay

Virgin OreR females were crossed with balancer line males (for chromosomes 2 and 3). Newly eclosed virgin F1 heterozygous females were collected and phenotyped for the presence/absence of the balancer (Figure S1). Following nutrient treatment, flies from both conditions were transferred to separate cages containing molasses plates. Wild-type males were added, and flies were allowed to mate for 24 hours. Eggs laid on each plate were counted, and plates were maintained in enclosed boxes with wet tissues to maintain humidity. After 48 hours, unhatched eggs were counted to calculate hatch rate and this procedure was repeated for 5 consecutive days.

Hatch rate is calculated as follows:

Hatch rate = (Total eggs laid - No. of unhatched eggs)/Total eggs laid

### Simulating meiotic crossovers in tetrads

A tetrad consists of two homologous chromosomes, each with two identical sisters. Chromosomal position was divided into 1000 bins. Bins are randomized in order and for each one, a Bernoulli draw was performed with probability *p* for CO to occur (i.e. CO rate per bin). Uniform across all bins, *p* = 2*d*/1000 where *d* is the length of the desired chromosome in Morgans when there is no drive. For every bin with a CO, 2 nonsisters were randomly selected to participate which causes reciprocal genotype exchange in all bins downstream. All simulations were run with 100,000 tetrads.

To implement interference, for every successful CO, we scaled the probability of CO placement for all remaining bins by

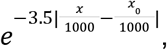

where *x* is the location of the bin and *x*_0_ the index of the bin where the interfering CO was placed. The constant − 3. 5 in the exponent was chosen such that the resulting simulated Genetic distance-to-RF curves approximated Kosambi’s mapping function. To prevent regional bias in which bins experienced interference, we attempted CO placement for each bin in a randomly shuffled order. Since our formulation of interference is reductional and decreases the number of placed COs, we increased the mean CO rate to recapitulate the same genetic distance after interference. To generate chromosomes with lengths of 1M, 1.5M, and 2M, the tetrad-wide CO-rate is 3.4, 7, and 13.78411 before interference which results in average COs-per-tetrad of 2, 3, and 4.

To implement asymmetric meiosis and recombinant drive, we created a pool of chromatids by selecting one from each tetrad. In the “loser-fall” regime, all chromatids with 1 or more COs were randomly selected for the pool. In the “winner-takes-all” regime, the chromatid with the most COs was always selected. E0 tetrads randomly transmit 1 chromatid. Strengths of drive were implemented by probabilistically choosing whether to use drive during chromatid selection for each chromatid.

Statistics such as the COs-per-chromatid distribution, interevent distance distribution, and morgans-to-RF curve were then tabulated from the resulting chromatid pool. To calculate CoC, we set a 75-bin “reference region” at the left end of the chromosome (bins 0-74), then set a “comparison region” from bins 74 to *x* and tabulated CoC between the reference and comparison regions for all *x*. To reduce noise, *x* was set starting from bin 149 (minimum comparison region size of 75 bins).

### Marker selected pool sequencing and processing

Adults were sexed and collected based on the presence and absence of the bw phenotype. Adult pools were processed by pulverization in liquid nitrogen and then DNA extraction as per Wei et al 2020 (36). WGS libraries were prepared using the Illumina DNA Prep Kit (Catalog # 20060059), QC’ed, and sequenced by Novogene on the Novaseq 10B 150PE configuration. Reads were aligned to the r6.68 reference with BWA (64), followed by genotyping in accordance with GATK’s (v4.4.0.0) best practice (65). Resulting vcf was filtered for diagnostic positions where the two parental strains (DGRP 360 and BL2336) are homozygous for different SNPs. Allele frequency at each diagnostic position was then inferred from the allele depth field of the vcf. For RF estimates, a modified LOESS non-parametric fit was used as per Wei et al. 2020 (36).

## Data availability

Raw reads have been deposited on the Sequenced Read Archive under PRJNA1521487.

