## Supplementary tables for "Conditions and properties of non-Mendelian transmission of recombinants in female meiosis"

Supplementary table S1. 4th chromosome recombinant data

| Markers | Treatment | Recombinants | Total | Recombinant frequency (RF) | p-value |
|---|---|---|---|---|---|
| *ci sv* | Nutrient-rich | 2 | 875 | 0.0023 | 0.83 |
| | Nutrient-deficient | 5 | 1326 | 0.0038 | |
| *ci sv ey* | Nutrient-rich | 0 | 241 | 0 | N/A |
| | Nutrient-deficient | 0 | 946 | 0 | |

Supplementary table S2. 2CO, interference, and CoC data

| Condition | Wild type strain | Markers | Treatment | 2CO (observed) | 2CO (expected) | CoC | Interference |
|---|---|---|---|---|---|---|---|
| 5-day deprivation | OreR | Chr 2: *al b c* | Standard | 114 | 96.2 | 1.19 | -0.19 |
| | | | Deficient | 123 | 129.18 | 0.95 | 0.05 |
| | | Chr 3: *hry ry e* | Standard | 64 | 79.67 | 0.80 | 0.20 |
| | | | Deficient | 56 | 78.38 | 0.71 | 0.29 |
| | | Chr X: *mtrm- w sn m* | Standard | 16 | 39.13 | 0.41 | 0.59 |
| | | | Deficient | 22 | 59.57 | 0.37 | 0.63 |
| | | Chr 2: *al dpy bw sp* (*al-bw*) | Standard | 39 | 44.45 | 0.88 | 0.12 |
| | | | Deficient | 51 | 67.39 | 0.76 | 0.24 |
| | | Chr 2: *al dpy bw sp* (*dpy-sp*) | Standard | 6 | 7.27 | 0.83 | 0.17 |
| | | | Deficient | 5 | 10.41 | 0.48 | 0.52 |
| | | Chr 2: *al dpy bw sp* (*al-sp*) | Standard | 4 | 2.49 | 1.61 | -0.61 |
| | | | Deficient | 0 | 3.64 | 0.00 | 1.00 |
| | | Chr 2: *al dpy bw sp* (3CO: *al-sp*) | Standard | 1 | 1.07 | 0.93 | 0.07 |
| | | | Deficient | 2 | 1.8 | 1.11 | -0.11 |
| | DGRP-360 | Chr 2: *al b c* | Standard | 53 | 72.14 | 0.73 | 0.27 |
| | | | Deficient | 101 | 124.96 | 0.81 | 0.19 |
| | | Chr 2: *al bw sp* (female) | Standard | 18 | 18.04 | 1.00 | 0.00 |
| | | | Deficient | 20 | 26.3 | 0.76 | 0.24 |
| | | Chr 2: *al bw sp* (male) | Standard | 24 | 27.36 | 0.88 | 0.12 |
| | | | Deficient | 23 | 26.99 | 0.85 | 0.15 |
| 12-days deprivation | OreR | Chr 2: *al b c* | Standard | 173 | 158.63 | 1.09 | -0.09 |
| | | | Deficient | 209 | 213.71 | 0.98 | 0.02 |
| | | Chr 3: *hry ry e* | Standard | 125 | 99.08 | 1.26 | -0.26 |
| | | | Deficient | 105 | 111.18 | 0.94 | 0.06 |
| | DGRP-360 | Chr 2: *al b c* | Standard | 94 | 97.15 | 0.97 | 0.03 |
| | | | Deficient | 168 | 186.19 | 0.90 | 0.10 |

Supplementary table S3. Fly strains information

| Strain type | Chromosome | Genotype | Stock Number | Source |
|---|---|---|---|---|
| Wild type | - | OreR | | Lab stock |
| | - | DGRP-360 | | Lab stock |
| Recessive mutant | X | *w sn m* | 147 | BDSC |
| | 2 | *al b c speck* | 210 | BDSC |
| | 2 | *al dpy bw speck* | 2336 | BDSC |
| | 3 | *hry ry e* | 93 | BDSC |
| | 3 | *yw; mtrm[exc13]/TM3sb* | 607084 | BDSC |
| | 4 | *ci sv* | 642 | BDSC |
| | 4 | *ci gvl ey sv* | 641 | BDSC |
| Balancer | 2 | *w; Sco/Cyo* | N/A | EG |
| | 3 | *yw; Dr/TM6c* | N/A | EG |

BDSC: Bloomington Drosophila Stock Centre

EG: Gift from Dr. Ethan Greenblatt
