## Supplementary figures for "Conditions and properties of non-Mendelian transmission of recombinants in female meiosis"

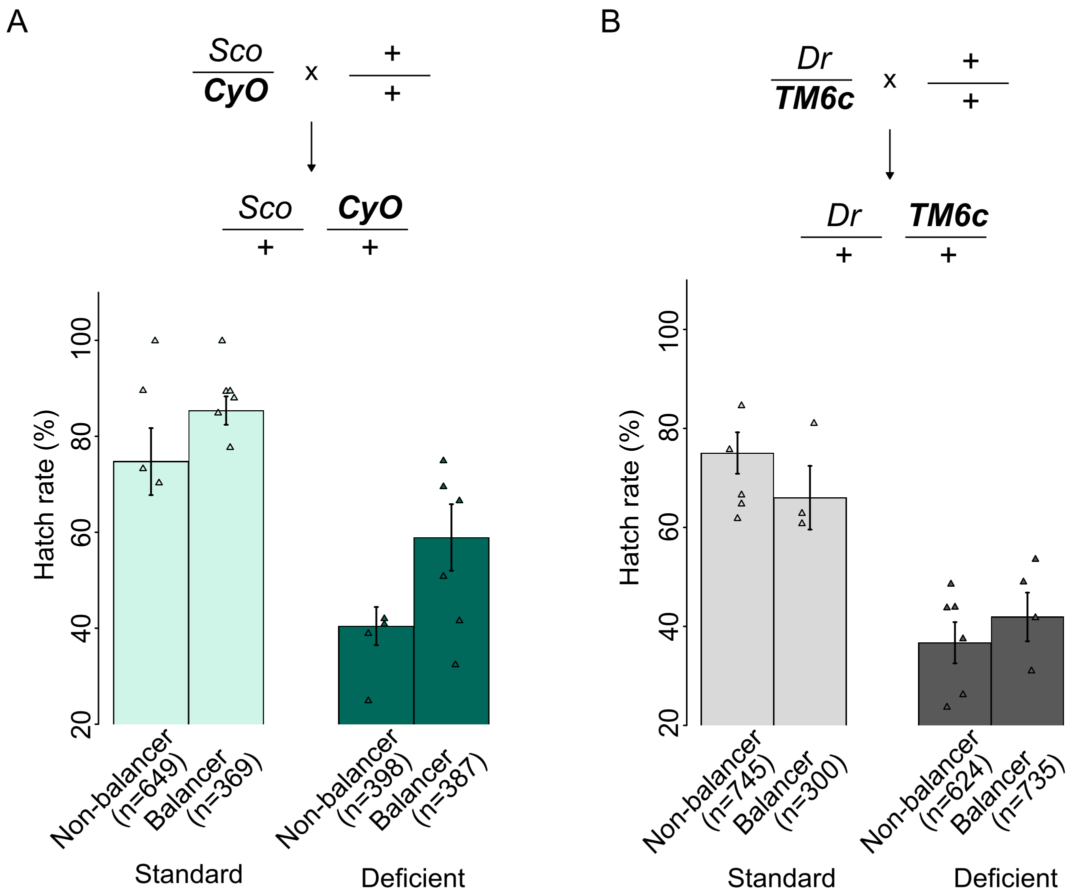


**Figure S1.** Hatch rate of chiasmate and balancer chromosomes. Top. Cross scheme to generate F1 females with and without 2nd (A) and 3rd (B) balancers. Bottom. Hatch rate estimates of progenies from females with and without balancers, subjected to standard or nutrient-deprivation conditions.

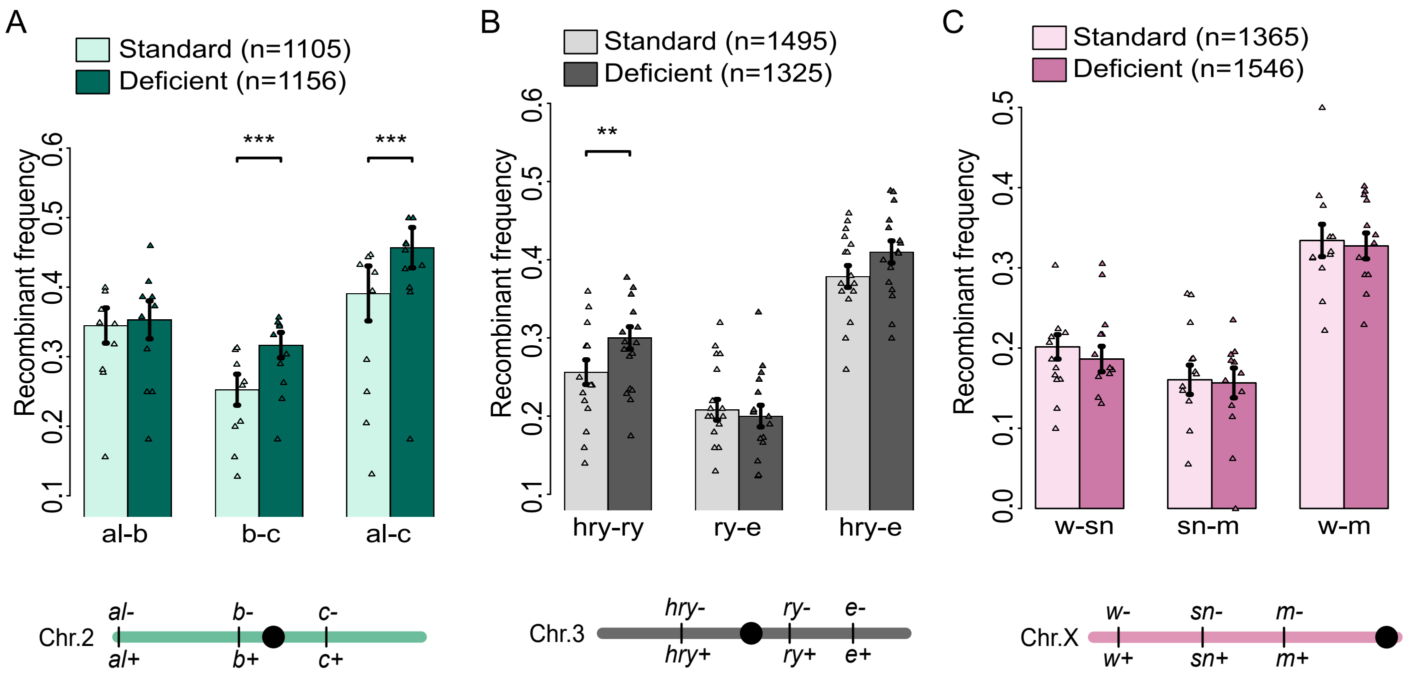


**Figure S2.** Recombination frequency increases after 5 days of nutrient-deprivation for Chr 2 (A), Chr 3 (B), and Chr X (C). *, **, and *** indicate p-values of <0.5, <0.05, and <0.005, respectively, inferred with the Chi-squared test.

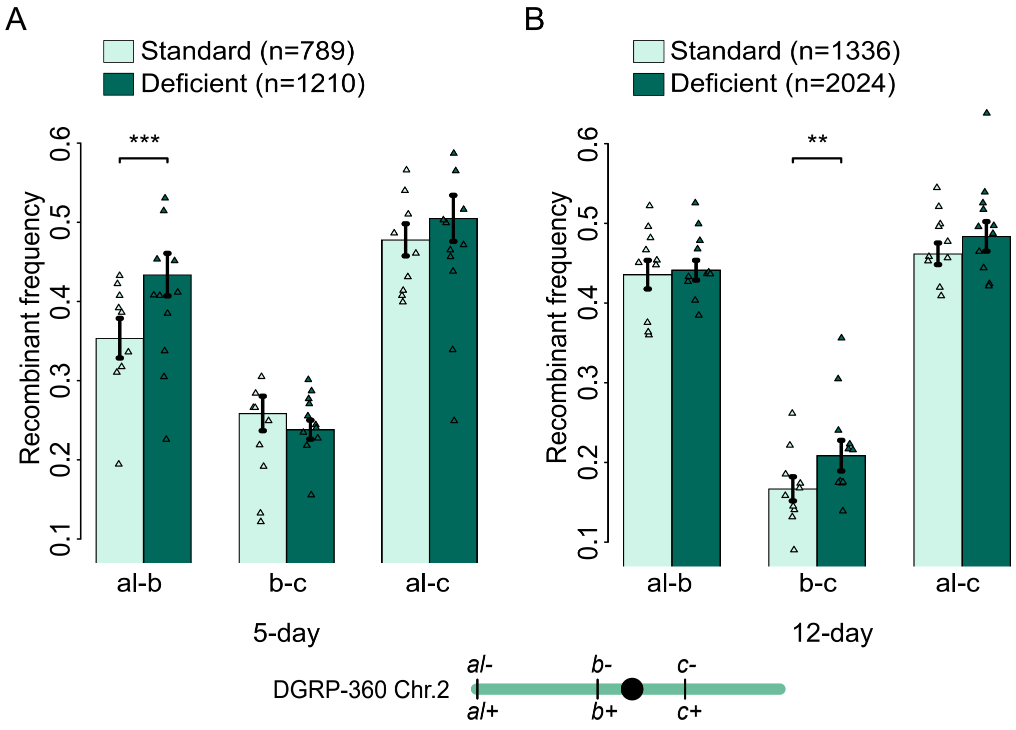


**Figure S3.** Recombination frequency on increases on Chr. 2 after 5 (A) and 12 (B) days of nutrient-deprivation in the DGRP-360 background.

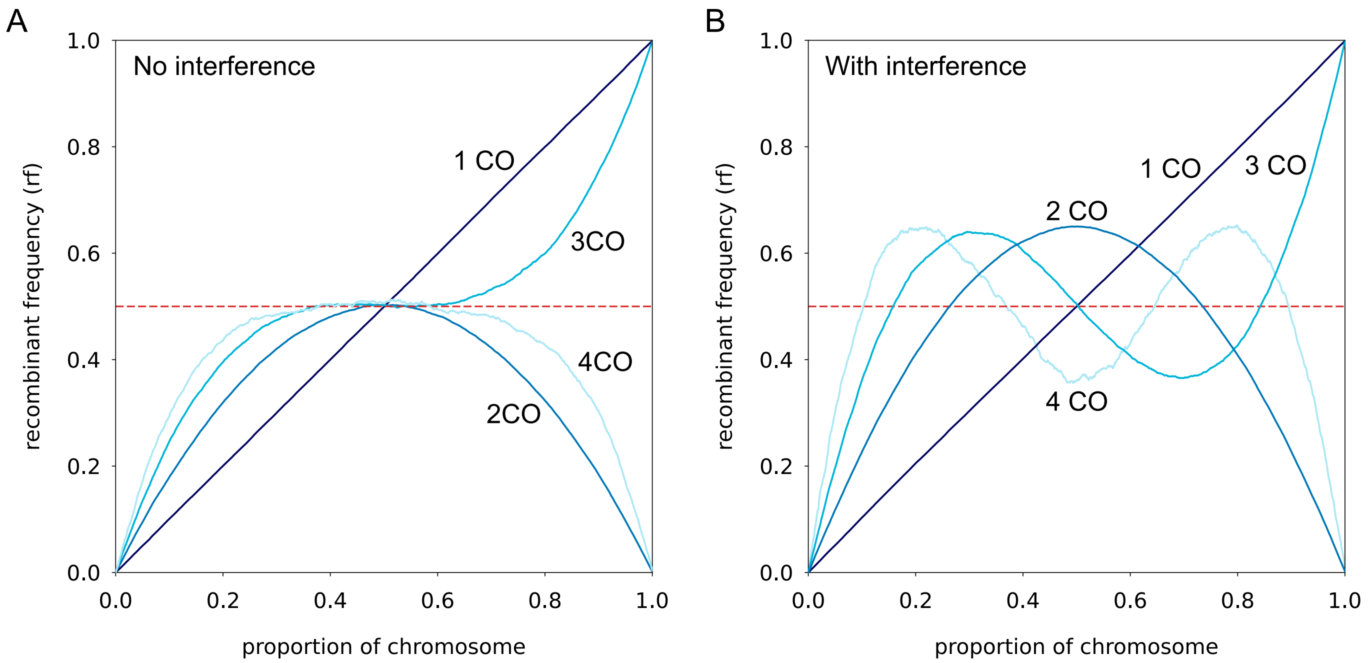


**Figure S4.** Behaviour of RF from chromatids different number of COs. COs are simulated with (A) and without interference (B).
